# Mis-splicing Drives Loss of Function of p53^E224D^ Point Mutation

**DOI:** 10.1101/2023.08.01.551439

**Authors:** Ian C. Lock, Nathan H. Leisenring, Warren Floyd, Eric S. Xu, Lixia Luo, Yan Ma, Erin C. Mansell, Diana M. Cardona, Chang-Lung Lee, David G. Kirsch

## Abstract

**Background:** Tp53 is the most commonly mutated gene in cancer. Canonical Tp53 DNA damage response pathways are well characterized and classically thought to underlie the tumor suppressive effect of Tp53. Challenging this dogma, mouse models have revealed that p53 driven apoptosis and cell cycle arrest are dispensable for tumor suppression. Here, we investigated the inverse context of a p53 mutation predicted to drive expression of canonical targets, but is detected in human cancer.

**Methods:** We established a novel mouse model with a single base pair mutation (GAG>GAC, p53^E221D^) in the DNA-Binding domain that has wild-type function in screening assays, but is paradoxically found in human cancer in Li-Fraumeni syndrome. Using mouse p53^E221D^ and the analogous human p53^E224D^ mutant, we evaluated expression, transcriptional activation, and tumor suppression in vitro and in vivo.

**Results:** Expression of human p53^E224D^ from cDNA translated to a fully functional p53 protein. However, p53^E221D/E221D^ RNA transcribed from the endogenous locus is mis-spliced resulting in nonsense mediated decay. Moreover, fibroblasts derived from p53^E221D/E221D^ mice do not express a detectable protein product. Mice homozygous for p53^E221D^ exhibited increased tumor penetrance and decreased life expectancy compared to p53 WT animals.

**Conclusions:** Mouse p53^E221D^ and human p53^E224D^ mutations lead to splice variation and a biologically relevant p53 loss of function in vitro and in vivo.

## Introduction

Tp53 is the most frequently altered gene in human cancer, as greater than 50% of cancers harbor p53 mutations. The primary p53 signaling axis is facilitated by protein stabilization through post-translational modification, homotetramerization and transcriptional upregulation of hundreds of target genes related to a diverse array of functions (1–3). Classically, the most critical functions of p53 for tumor suppression were thought to be the canonical pathways leading to cell cycle arrest, apoptosis and senescence. Challenging this model, a triple knockout mouse with deletion of three critical p53 targets driving cell cycle arrest, Cdkn1a, and apoptosis, Puma and Noxa, did not exhibit a significant tumor burden (4–6). This suggested that these effectors are insufficient or potentially unnecessary for p53 mediated tumor suppression in mice. In support of the insufficiency of these canonical pathways of tumor suppression, mice with p53 mutations in the transactivation domain (p53^25,26^) or in three lysines (p53^K117R+K161R+K162R^), which lack canonical signaling to initiate cell cycle arrest and apoptosis after DNA damage nevertheless retained tumor suppression (7,8).

In human cancers, 86% of mutations in p53 occur in the DNA binding domain and ∼30% of these mutations are specific to the top six hotspot residues (9,10). The impact of these hotspot mutations on the structure and function of p53 is well characterized. These mutations are classified as structural mutations that significantly affect the overall architecture and stability of the DNA binding domain or contact mutations that alter residues critical for DNA interaction (11). Hotspot mutants are further characterized by their accumulation at the protein level due to disrupted negative feedback loops with potential neomorphic capabilities (12). These mutations also lead to loss of signaling through critical effectors of p53 function, a phenotype confirmed in mouse models (12–15). For example, the p53^R175P^ and p53^E177R^ mouse models retain the ability to drive cell cycle arrest, but exhibit a loss of canonical p53 mediated apoptosis. The p53^R175P/R175P^ and the p53^E177R/E177R^ mice display a delayed onset of tumorigenesis relative to p53^-/-^ (16,17). Therefore, these mouse models of a DNA binding domain mutants only partially discriminate canonical signaling from tumor suppression.

Systematic characterization of the functional impact of point mutations in p53 on transactivation was first explored in yeast (18). Investigators generated single base pair amino acid substitutions in the previously uncharacterized portion of the DNA binding domain as well as the NH_2_- and COOH-terminal domains. Using these mutants, investigators examined differential transactivation of eight critical effectors of p53 function (18). As expected, loss of function mutations clustered in the DNA binding domain and the transactivation domain. This confirmed previous reports that other functional domains are insensitive to single amino acid substitutions (19–23). Systematic studies in mammalian cell lines confirmed a distribution of transactivation functionality of p53 mutants (14,24).

To explore the link between transactivation and tumor suppression, here we compare the findings from the yeast screen study with available human p53 mutation data from cancers to correlate specific point mutations showing loss or retained transactivation function with the incidence of the same substitution in human cancer. We identified a subset of mutants that included p53^E224D^ as competent for transactivation in yeast, but paradoxically present in human cancers. Because the ability of a mutant of p53 to suppress cancer has historically been tightly linked with a specific capacity to drive the expression of transcriptional targets, we characterized the transactivation potential of cDNAs of these mutants in human cells and confirmed that p53^E224D^ retained transactivation. Therefore, we generated genetically engineered mice with an endogenous p53^E221D^ mutation (the mouse equivalent of p53^E224D^). Using in-vitro experiments with mouse embryonic fibroblasts (MEFs) and human cancer cells harboring p53^E224D^, and the in-vivo genetically engineered mice, we find that the p53^E224D^ point mutation loses the ability to suppress tumor development because it causes a loss of p53 function as a result of mis-splicing of the p53 mRNA transcript.

## Results

### Transactivation Mutants Co-Occur with Secondary DNA Binding Domain Mutations

To search for p53 mutants with intact transactivation in the yeast model system and present in human cancer, we searched the IARC, TCGA, and GENIE tumor databases for the relative frequency of p53 mutations in patient tumor samples (Figure 1A). The yeast screening data identified 373 mutants classified as competent or wild-type for transactivation and simultaneously found in human tumor databases of p53 mutations (18). Co-occurrences, instances of a transactivation competent mutant in a sample with a secondary non-synonymous p53 mutation, are noted because the overall loss of tumor suppressive capacity could potentially be attributed to either mutation if one mutant had dominant negative activity. Of note, the p53^E224D^ mutation occurs frequently compared to other mutants without a substantial number of co-occurring non-synonymous p53 mutations. In addition, the p53^E224D^ mutation was also found in the IARC database among germline, transactivation competent, mutants in the p53 gene (Figure 1B). These results suggest that p53^E224D^ is a primary event driving altered function of p53 in these tumors.

**Figure 1:**
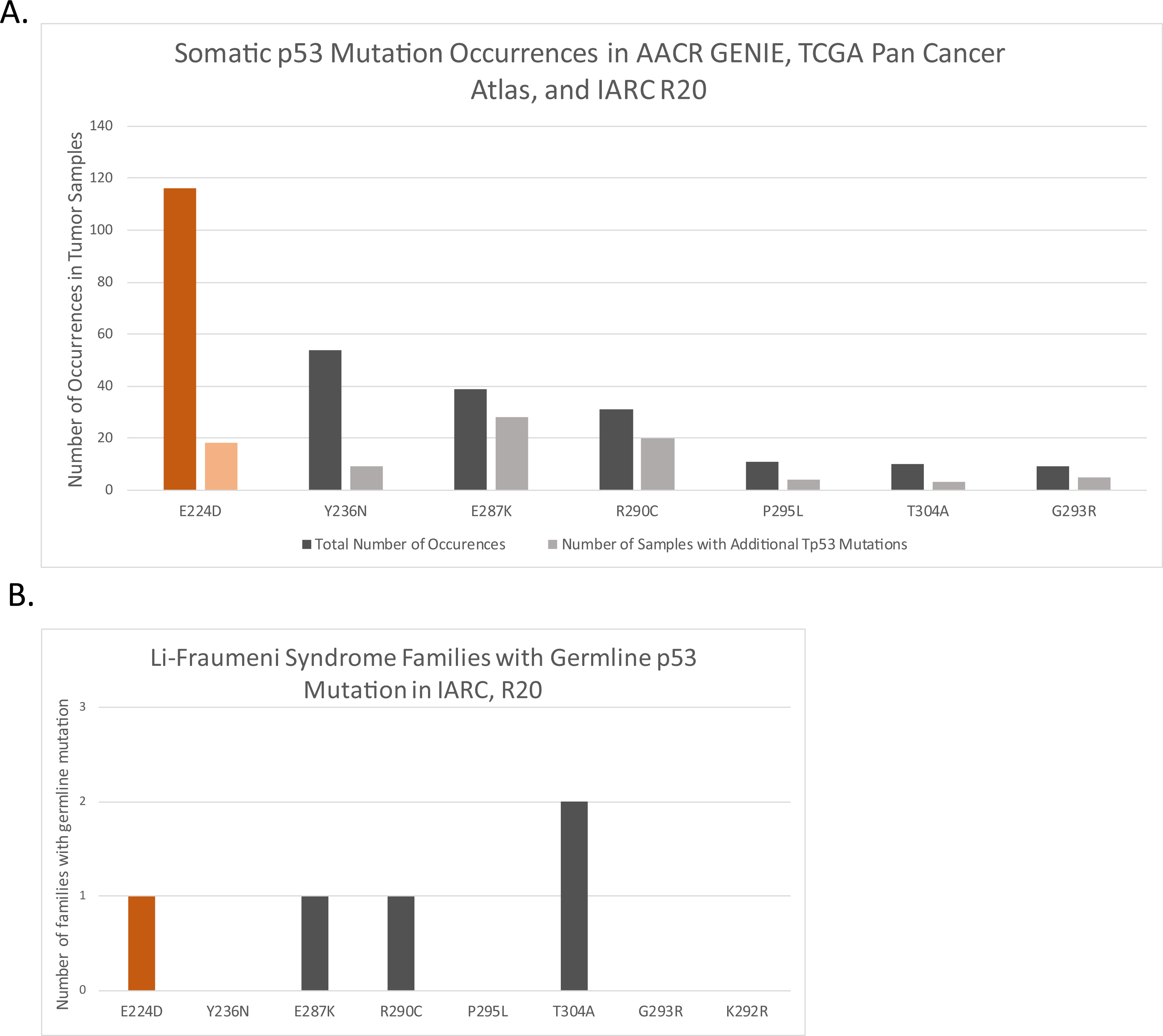
Incidence of p53 transactivation competent mutants in human tumor data. A. Somatic occurrences of select transactivation competent p53 mutants from IARC, GENIE, and TCGA databases. Lighter adjacent bars represent the co-occurrence of non-synonymous p53 mutants in the same samples. B. Germline occurrences of p53 mutants from IARC database

### p53^E224D^ cDNA Retains Transactivation in Human Cells

To evaluate the relationship of the yeast transactivation screen within a mammalian system, we used a tetracycline-inducible system to express various p53 mutants in the p53 null non-small cell lung cancer cell line H1299. H1299 cells have a homozygous deletion of a portion of the p53 gene and no protein expression to interfere with this re-expression assay (25). We transduced the H1299 cells with a tet-inducible backbone containing a doxycycline responsive cDNA that encodes p53 wild-type (WT) or various mutants identified in the yeast screen. Doxycycline addition was sufficient to overexpress p53 wild-type (WT) or mutant proteins and to assess their impacts on p53 targets MDM2 and CDKN1A by western blot (Figure 2A). In this assay, human p53^E224D^ expressed from cDNA retained the capacity to activate the key p53 target proteins MDM2 and CDKN1A.

**Figure 2:**
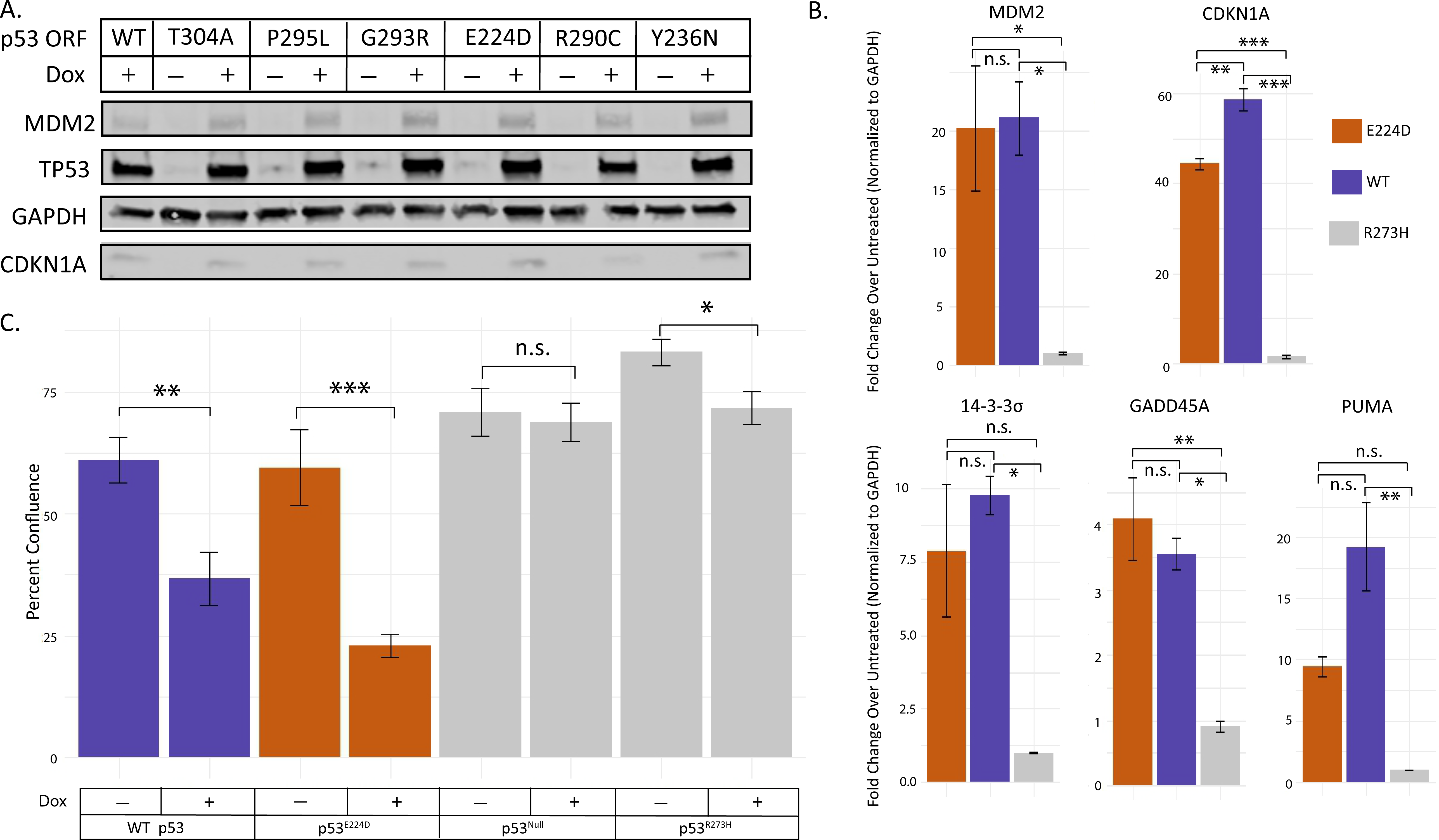
Capacity of Selected p53 Mutants for Transactivation. A. Expression of doxycycline inducible p53 mutant cDNA constructs in p53 null H1299 lung cancer cells with expression validated by western blot using cell lysates probed for p53. Transactivation capacity assessed by MDM2 and CDKN1A expression using a GAPDH loading control 24 hours post doxycycline addition, 1µg/mL. B. Transactivation of p53 targets by doxycycline (24 hours, 1µg/mL) induced p53^E224D^ mutant 1 hour after 10 Gy irradiation. Gene expression assessed by RT-qPCR. (p53^E224D^ – Orange, WT – Purple, p53 Loss of Function (R273H) – Gray) (* = p<.05) Average of three technical replicates. C. Incucyte measurement of growth arrest capacity of p53 mutants. Percent confluence of samples represented as an average after six days of growth in a 96 well plate. Compared to non-doxycycline containing media over the same time course (p53^E224D^ – Orange, WT – Purple, p53 Loss of Function (Null or R273H) – Gray) (*** = p<.001, ** = p<.01, * = p<.05) Average of ten technical replicates.

The transactivation capacity of the p53^E224D^ mutant was further assessed using qRTPCR after doxycycline and irradiation treatments to induce a p53 transcriptional program. As expected, transactivation from the doxycycline induced p53^E224D^ mutant was not significantly different from p53 WT for key transcriptional targets of p53 that regulate apoptosis (PUMA), cell-cycle arrest, and senescence (CDKN1A, 14-3-3 sigma, and GADD45A) as well as the E3 ubiqitin ligase that degrades p53 protein (MDM2) (Figure 2B). In contrast, the expression of these p53 target genes was not induced by a dominant negative R273H mutant (Figure 2B). To test if transactivation of these p53 targets by p53^E224D^ leads to cellular phenotypes, we compared the cell growth in H1299 cells harboring doxycycline-inducible p53^WT^, p53^E224D^, or p53^R273H^ as well as the p53 null parental cells in the presence of doxycycline containing or control media. Growth was tracked via Incucyte over a six-day period. Cumulative growth assessed on the final day is reported as average percent confluence (Figure 2C). p53 WT and p53^E224D^ mutants induced by doxycycline retained growth suppression. Together, our results indicate that the cDNA for p53^E224D^ retains the ability to transactivate downstream targets and suppress the growth of human cancer cells in vitro.

### A p53^E221D^ genetically engineered mouse model lacks p53 protein and RNA expression

Although the p53^E224D^ mutation does not significantly impair the function of p53 protein in the setting of cDNA expression, the impacts of the p53^E224D^ mutation on the expression and function of p53 protein from the endogenous p53 locus in-vivo remained unclear. Thus, we generated a novel genetically engineered mouse with a p53^E221D^ mutation (GAG>GAT, mouse equivalent of human p53^E224D^) in the endogenous p53 gene through homology-directed CRISPR editing. The presence of the p53^E221D^ mutation was validated using targeted sequencing of the p53 gene (Supplementary Figure 1A-C). We generated mouse embryonic fibroblasts (MEFs) from the p53^E221D^ mice. Unexpectedly, we observed that p53 protein was not detected in p53^E221D/E221D^ MEFs from the knock-in mice even after treatment with either 10Gy irradiation or with the proteasomal degradation inhibitor MG132 (Figure 3A). Similar to the results from our mouse model, the human colorectal cancer cell line NCI-H716 harboring a homozygous p53^E224D^ mutant also showed no detectable p53 protein after treatment (Figure 3B). Consistent with a loss of p53 function, we observed that both p53^E221D/E221D^ and p53^E221D/WT^ mice showed a significant decrease in overall survival compared to p53^WT/WT^ littermates (Figure 3C). These mice primarily developed sarcomas and thymic lymphomas, which are also the predominant types of tumors that occur in mice with germline deletion of p53 (Supplementary Figure 2A) (26–28). The mean survival of p53^E221D/E221D^ is 167 ± 57 days, which is similar to the mean survival of p53^-/-^ mice (27,29).

**Figure 3:**
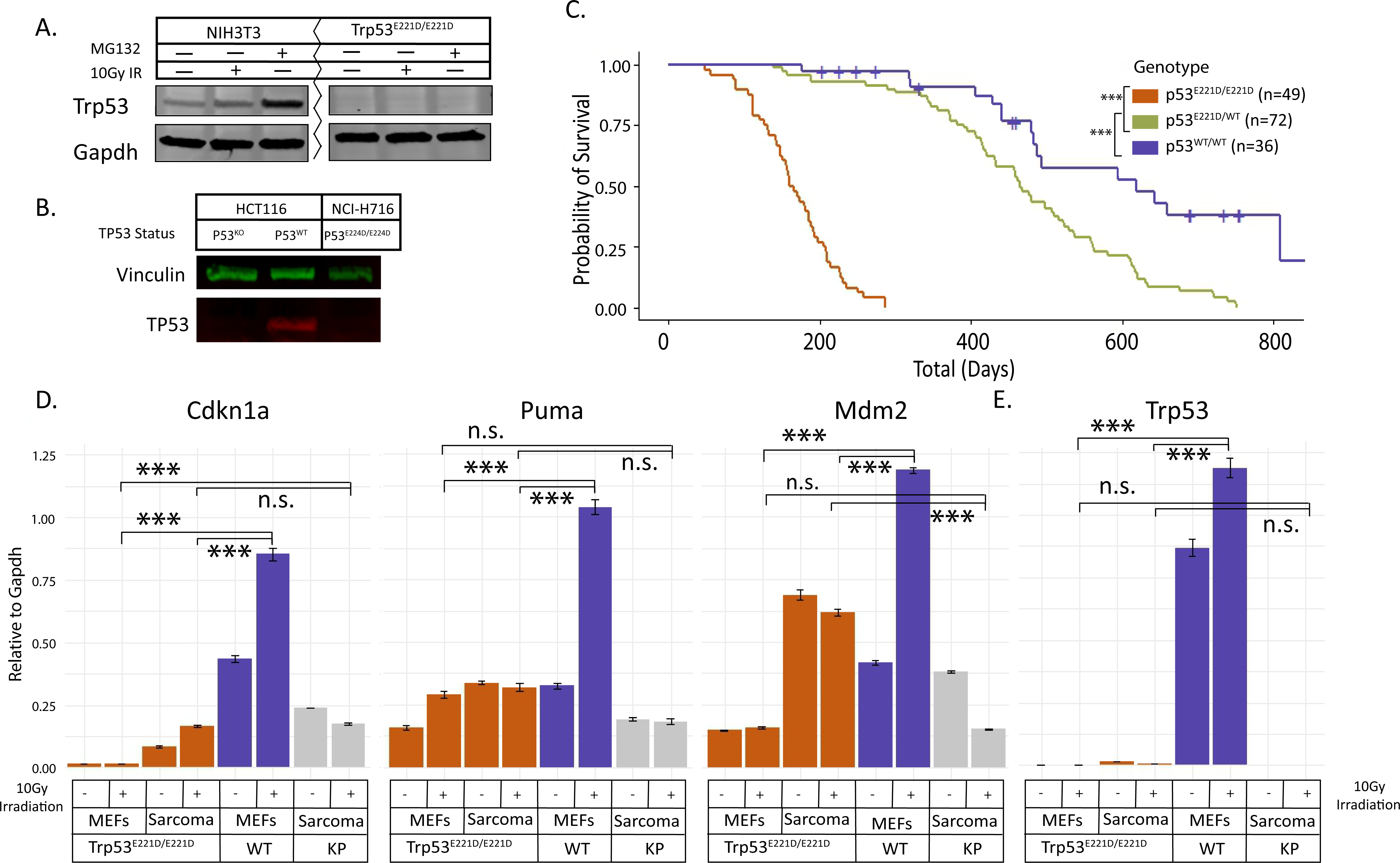
In-Vivo Function of the mouse p53 P53^E221D/E221D^, which is analogous to human p53^E224D^. A. Mouse embryonic fibroblast (MEF) lysates were probed via western blot after stabilization hour after 10 Gy irradiation or 6 hours after addition of the proteasomal degradation inhibitor MG132, 10µM. B. Human colorectal adenocarcinoma cell line, NCI-H716 which harbors the p53^E224D^ mutation, and p53^WT^ and p53^KO^ human colorectal carcinoma, HCT116, probed for p53 protein after MG132 (6 hours, 10µg), G418 (48 hours, 400µg/mL), and Irradiation (1hr before extraction, 10Gy) treatment. C. Survival Curves for p53 mutant mice. Mice monitored until ill health necessitated euthanasia or found dead. (p53^E221D/E221D^ – Orange, p53^WT/WT^ – Purple, p53^E221D/WT^ – Green) (*** = p<.001 using pairwise log-rank test) D. p53 target gene expression changes, assessed by RT-qPCR, one hour after 10 Gy irradiation. (p53^E221D/E221D^ – Orange, p53^WT/WT^ – Purple, p53^-/-^ Kras^G12D^ - Gray) (*** = p<.001, ** = p<.01, * = p<.05) Average of three technical replicates.

Compared to p53^WT/WT^ MEFs, the induction of transcriptional targets of p53 by ionizing radiation was markedly impaired in homozygous p53^E1221D/E221D^ MEFs or in a cell line derived from a homozygous p53^E221D/E221D^ sarcoma (Figure 3D). Of note, the expression of p53 mRNA was absent in p53^E221D/E221D^ MEFs (Figure 3E). Collectively, our results from p53^E221D/E221D^ mice indicate that p53^E221D^ is a loss-of-function mutation that results from impaired expression of endogenous p53 mRNA in vivo. These findings generated a hypothesis that mis-splicing of the p53^E221D^ mutant leads to nonsense mediated decay as a potential mechanism for this loss of function.

### Mis-splicing drives loss of RNA expression

To investigate potential mis-splicing from the p53^E221D^ mutation, RNA-Seq was performed on p53^E221D/E221D^ MEFs and the p53^E224D/E224D^ human cell line. Transcript analysis of TopHat aligned RNA sequencing from homozygous p53^E221D/E221D^ and p53^WT^ MEFs supports a general loss of p53 transcripts at the RNA level (Figure 4A). Using stringtie, a bioinformatics tool for splice variation, four novel transcripts that include the exon 6 splice junction were identified in the mutant MEFs. Novel transcripts MSTRG.3218.1, MSTRG.3218.5, and MSTRG.3218.7 highlight exon skipping, and MSTRG.3218.8 retains intron 6 (Figure 4A). The scale of the Sashimi plots from the Integrative Genomics Viewer (IGV) visualization of a representative sample confirms the reduction in read depth at this region of the p53^E221D/E221D^ mRNA. This plot also allows visualization of the 43 reads that confirm splicing of the Exon 5 donor to the Exon 7 splice acceptor and the relative increase in reads that map to Intron 6 in the p53^E221D/E221D^ MEFs. (Figure 4B). Of note, a small number of reads in each sample are correctly spliced, one in the representative sashimi visualization. These are visual confirmations of the transcript analysis that discovered novel transcripts with complete intron 6 retention and exon 6 skipping. Transcript discovery was subsequently carried out using Star alignments (Supplementary Figure 3A-B). Transcript structure was broadly confirmed, but there was variability in the mutant transcript prevalence. The wild-type Trp53 transcripts were well conserved across alignment methods.

**Figure 4:**
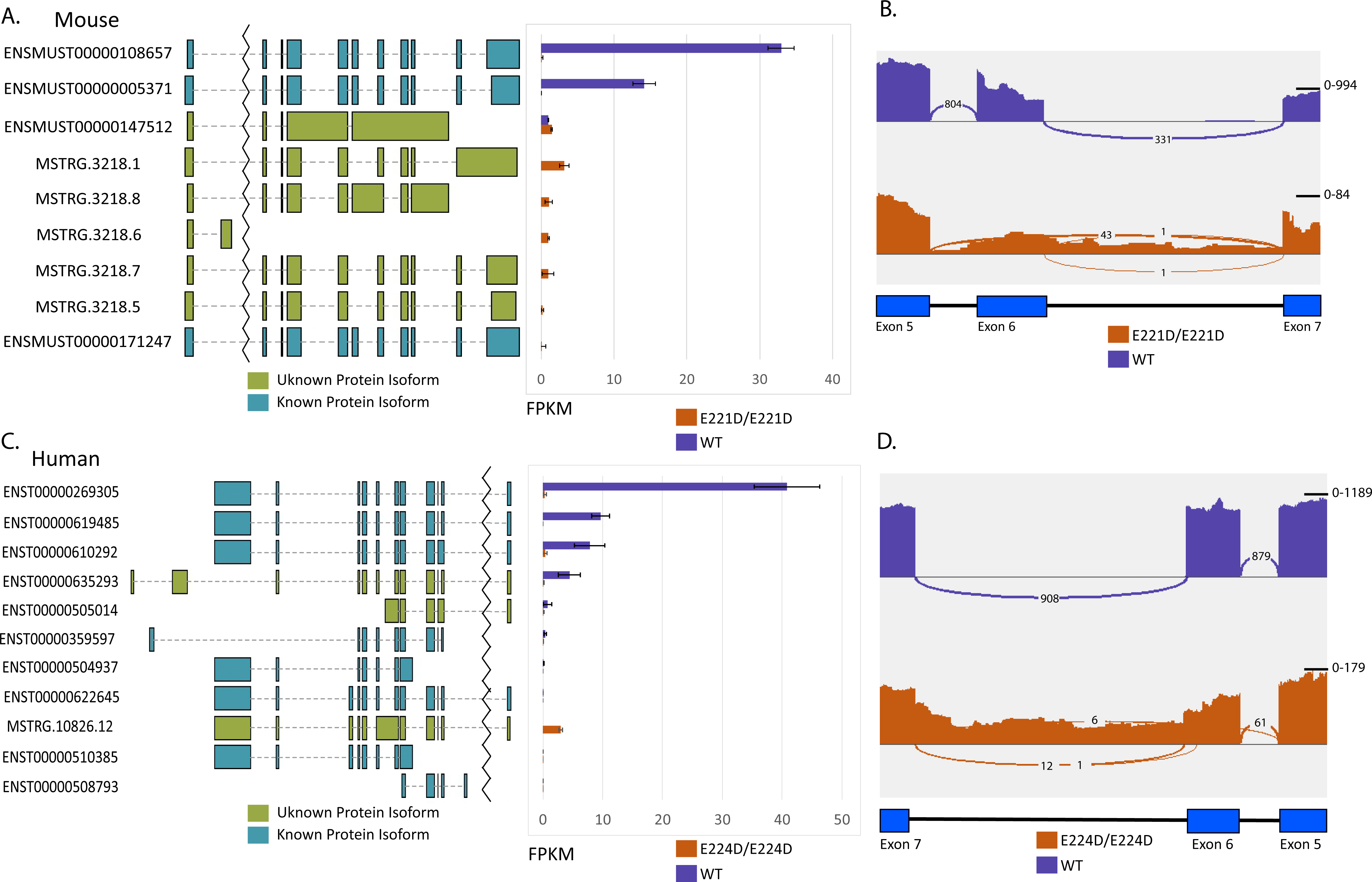
Transcript Identification in p53 Mutant Samples. A. Visual reference for RNA transcript splicing adjacent to bar plot of relative abundance (FPKM) in p53^E221D/E221D^ and p53^WT^ MEFs. Known transcripts annotated with Ensembl identifier, unknown transcripts annotated with unique Stringtie identifier. Color of the graphical representation of RNA transcript symbolizes predicted translation of protein of a known or unknown function. Sequencing done on three technical replicates. B. Representative sashimi plot visualization of read abundance from IGV. Linkages represent reads that span exon-exon junctions, i.e. splice events in MEFs. C. Visual reference for RNA transcript splicing adjacent to bar plot of relative abundance (FPKM) in p53^E224D/E224D^ human cancer cell line NCI-H716 and p53^WT^ cancer cell line HCT116. Known transcripts annotated with Ensembl identifier. Color of the graphical representation of RNA transcript symbolizes predicted translation of protein of a known or unknown function. Sequencing done on three technical replicates. D. Representative sashimi plot visualization of read abundance from IGV. Linkages represent reads that span exon-exon junctions, i.e. splice events for human cell lines.

In contrast to the Tophat aligned novel transcript identification in the mouse model, when we performed RNAseq on human p53^E224D/E224D^ NCI-H716 compared to p53^WT^ HCT116 cells, there was one novel variant highlighted after Stringtie transcript identification (Figure 4C). This transcript variant, MSTRG.10826.12, exhibits intron 6 retention as the dominant transcript in the p53^E224D/E224D^ cell line (Figure 4D). In the Star aligned reads, however, Stringtie results suggest complete intron retention as the dominant transcript while a second novel transcript results from splicing at a cryptic splice site 5 base pairs downstream of the mutation. (Supplementary Figure 3E). Similar to the sequencing results in the MEFs, wild-type Tp53 transcripts are well conserved across sequence alignment methods. Together, these results show that the p53^E221D^ mutation in mice and the p53^E224D^ mutation in humans results in the loss of RNA expression through mis-splicing.

### Inhibition of nonsense mediated decay does not significantly alter transcript composition

One pharmacological strategy to rescue prematurely degraded p53 RNA is to inhibit nonsense mediated decay and simultaneously encourage ribosomes to read through a premature stop codon. This could potentially result in the expression of a functional p53 protein that restores the ability of cells to activate p53-mediated tumor-suppressive pathways (30–33). To assess the p53^E221D^ mutant as a candidate for this treatment combination, cells were treated with G418 to promote ribosome readthrough and the SMG7 inhibitor NMDI14 to inhibit non-sense mediated decay. We hypothesized that artificial increases in the transcript levels could increase correctly spliced transcripts to a physiologically relevant level. Using the combination described above, p53 RNA levels were significantly increased in both MEFs and a tumor derived sarcoma cell line from p53^E221D/E221D^ mice, however full length p53 RNA transcripts were not generated (Figure 5A-D).

**Figure 5:**
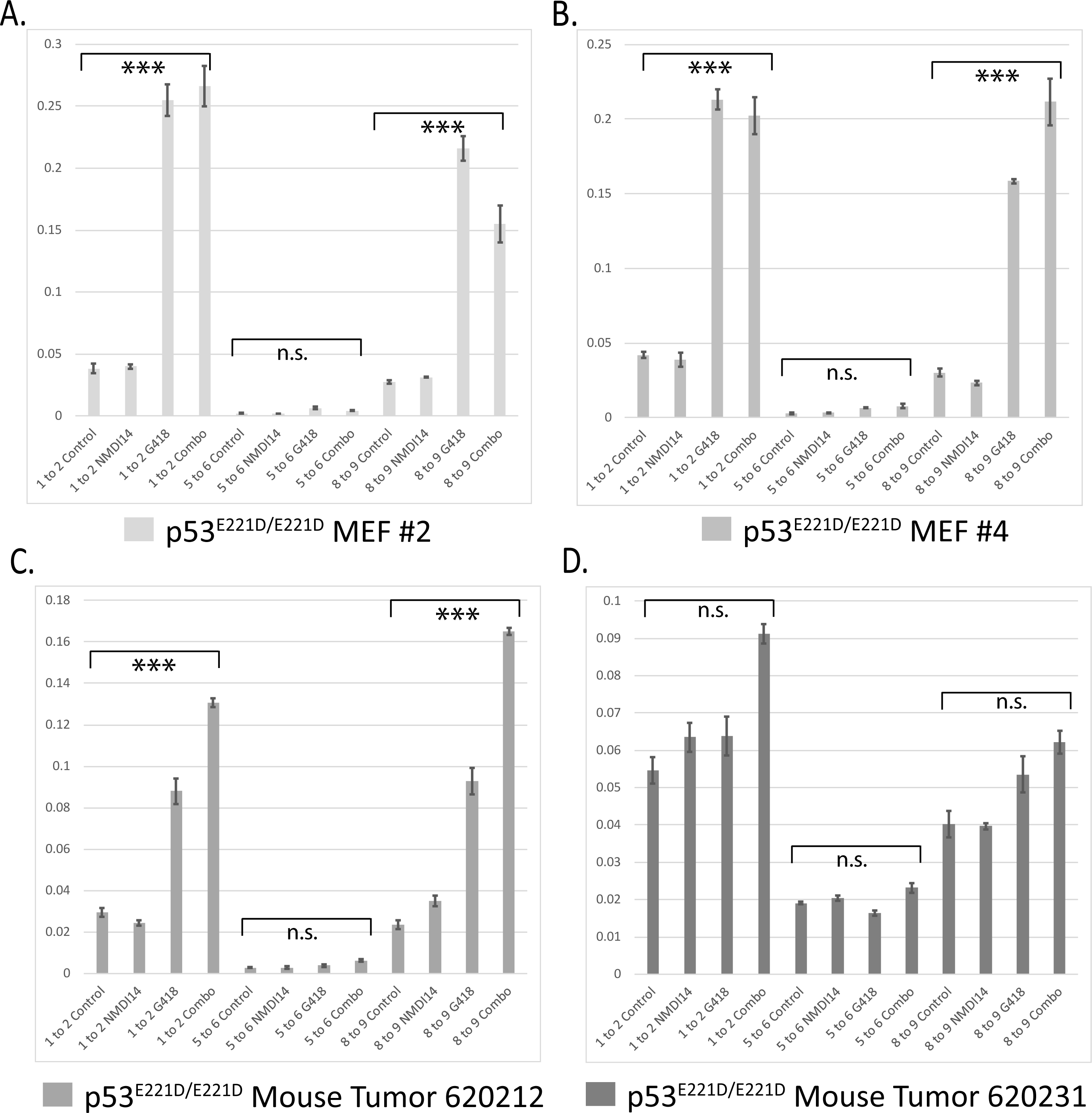
Nonsense Mediated Decay Inhibition in a Mouse Model. RNA was harvested from MEFs and mouse tumors homozygous for the P53^E221D/E221D^ mutation. Probes spanning the p53 Exon1/Exon2, Exon 5/Exon6, and the Exon8/Exon9 junctions were used to assess any changes in the usage of these exons after treatment with G418, 400µg/mL, and the SMG7 inhibitor NMDI14, 5µM, for 48 hours. A. MEF cell line #2, p53^E221D/E221D^ Average of three technical replicates. B. MEF cell line #4, p53^E221D/E221D^ Average of three technical replicates. C. Sarcoma cell line from p53^E221D/E221D^ mouse 620212 Average of three technical replicates. D. Sarcoma cell line from p53^E221D/E221D^ mouse 620231 Average of three technical replicates.

## Discussion

p53 DNA binding missense mutants continue to be an expanding area of research for designing new therapies targeted towards stabilization and reactivation of the proteins diverse critical pathways for tumor suppression (Reviewed in (34–37)). However, the impact of p53 mutations that lie outside of the most frequently occurring hotspot regions when expressed from the endogenous promoter in multiple cell lineages remains less well-investigated. In this study, we sought to understand the tumor suppressive dynamics of point mutations in the DNA binding domain of p53 that paradoxically retain transactivation of canonical targets, but are still found in human cancer. We narrowed a list of candidate mutations from a published screen in yeast where the mutants showed approximately wild-type transactivation for eight select promoters of p53 target genes and where the mutations are also found in human cancer. We used a human p53 null cancer cell line to validate transactivation of the cDNA of the p53^E224D^ in a mammalian system and selected the p53^E224D^ mutant for further study because it had been found in human cancers and in Li-Fraumeni syndrome. Therefore, we generated a genetically engineered mouse model with the analogous p53^E221D^ mutation. Contrary to our expectation, however, the mouse model harboring the p53^E221D^ mutant did not express detectable p53 protein. This was further observed in a human p53^E224D/E224D^ cancer cell line. Low p53 RNA levels in both the human and the mouse cell lines suggested that mis-splicing drives nonsense mediated decay of the mutant p53 mRNA.

Our findings in the human samples are consistent with previous studies that did not observe detectable p53 protein in the NCI-H716 cell line as well as experiments that used available human data and in-vitro assays to investigate alternative splicing at this site (38–41).These studies found partial intron retention at the 5’ end of intron 6 was present after both the synonymous G>A mutation or the G>T/C mutations for p53^E224D^ as we observed (38–44).

Our study contributes an expanded understanding of the intron retention in p53^E224D^ in human cells as well as developing a novel p53^E221D^ mouse model of mis-splicing of p53. Our sequencing suggests that there are multiple transcripts present following mis-splicing at this site. Consistent with previous studies on human data, the first 5 base pairs of intron 6 are retained in one transcript while larger portions of intron 6 are retained in other transcripts. To our knowledge, the p53^E221D^ variant that was generated in this study is the first mis-spliced variant of Trp53 to be explored in a mouse model. The variation in splicing that we observed in differential cryptic splice site usage and exon skipping provides a mouse model to investigate approaches to reactivate p53 by targeting nonsense mediated decay. Moreover, this model functions as a p53 knockout model that can be used as a more biologically relevant model in studies investigating point mutant variability in the DNA binding domain.

## Materials and Methods

### Database Access

IARC version R20 accessed at (https://tp53.isb-cgc.org/), TCGA (https://www.cbioportal.org/), and GENIE (https://genie.cbioportal.org/) database accessed for non-redundant studies with human P53 mutational data.

### Mouse Model Generation and Animal Use

P53^E221D^ mice were generated by sgRNA, CAS9 (IDT cat# 1081059), and single stranded oligodeoxynucleotide, ssODN, electroporation of mouse embryo as previously described (45). Single guide RNA was synthesized via in-vitro transcription following the protocol described previously (Supplementary Table 1) (45). Repair single stranded oligodeoxynucleotide repair oligo was purchased from Integrated DNA Technologies (Supplementary Table 1). This process generated a cut site in the p53 locus at the desired mutation region and repair was guided using a homologous template for repair with the intended mutation (Supplementary Figure 1A). Chimeric mice were tested for successful targeting of the vector into the p53 locus and confirmed via PCR (Supplementary Table 1). PCR products subcloned into pL253 via in-fusion cloning and sequenced for target validation.

Mice were monitored weekly when possible (restricted during COVID19 lockdown procedures). Upon mortality or humane endpoint, tumors harvested and saved for histology and flash frozen if tumor size and condition allowed. All animal studies were performed in accordance with protocols approved by the Duke University Institutional Animal Care and Use Committee.

### Genotyping

Mice were genotyped using tails collected from mouse pups. For early mouse generations tail genomic DNA was isolated and PCR was performed using primers listed in Supplementary Table 1. PCR products were visualized after electrophoresis in 1.5% agarose gels. Subsequent generations, genotyping was carried out by Gene Master LLC.

### Immunohistochemistry

Tumors were fixed in formalin overnight, transferred to 70% ethanol, paraffin-embedded, and sectioned to 5 μm thickness and stained with H&E. Tissue sections were examined by a sarcoma pathologist (D.M. Cardona).

### Cell Lines and Reagents

HCT116 WT and HCT116 KO cells were a gift from Bert Vogelstein. NCI-716 and H1299 cells were obtained from ATCC and grown in DMEM media (Gibco), 10% FBS (Avantor), and 1X Antibiotic-Antimycotic (Gibco). H1299 cells expressing doxycycline inducible p53 WT and mutant constructs were generated by lentiviral infection in the Duke Functional Genomics Facility. Cells were cultured in 2µg/mL doxycycline for 24 hours before WB and qRTPCR experiments. Primary mouse embryonic fibroblasts were isolated from E12.5 to 13.5 mouse embryos and cultured in 37°C, 5% CO_2_ conditions. Cells were treated with G418 at 400µg/mL (Geneticin™ for 48 hours, ThermoFisher), NMDI14 was used at 5µM (Calbiochem) for 48 hours, and MG132 at 10µM (SelleckChem) for 6 hours. Cells were harvested following 1 hour following 10 Gy irradiation. Vehicles for G418 (water), NMDI14 (DMSO), and MG132 (DMSO) added to control conditions.

### Radiation Treatments

Cells were plated at least 24 hours prior to irradiation. The X-RAD 160 (Precision X-ray) cell irradiation system with 2 mm aluminum filter was used to deliver 10 Gy to cells. The Duke University Radiation Safety Division staff maintains and performs dosimetry for the irradiator.

### Lentivirus Production

All lentiviruses for Tet-inducible expression of p53 mutants were prepared by cotransfecting HEK293T cells with the lentivector, psPAX2 (Addgene #12260) and pMD2.g (Addgen #122259) using TransIT-LT1 (Mirus) according to manufacturer instructions. Supernatant containing lentivirus was harvested at 48 hours, aliquoted and stored at -80 for transductions. H1299 cells were transduced with p53 mutant lentivirus as indicated at a multiplicity of infection (MOI) of <1 in the presence of 4µg/ml polybrene. Twenty-four hours post transduction, cells were selected with 2µg/ml puromycin for 72 hours and then expanded for use in experiments.

### Immunoblots

Cells were lysed with RIPA (Thermo Scientific) supplemented with PhosStop Tablet (Roche), Aprotinin (Sigma Aldrich), Halt Protease and Phosphatase Inhibitor Cocktail and further lysis achieved by 10 second sonication (QSonica). 30µg of protein was subjected to SDS-PAGE and wet transferred to nitrocellulose membranes. Blots were blocked using Interceptor Blocking Buffer (LI-COR) for a minimum of one hour before probing with primary antibodies. Immunoblots were probed with antibodies against mouse p53 (Cell Signaling, 1C12), Gapdh (ProteinTech, 60004-1) human p53 (Invitrogen, PA5-27822), MDM2 (Santa Cruz Biotechnology, sc-965), and CDKN1A (Santa Cruz Biotechnology, sc-6246). Anti-rabbit IRDye® Secondary Antibodies, RDye® 680RD Donkey anti-Mouse IgG Secondary Antibody (LI-COR, 926-68072), or IRDye® 800CW Goat anti-Rabbit IgG Secondary Antibody (LI-COR, 926-32211) were used. The signal was detected using Odyssey imaging system (LI-COR Biosciences).

### RNA Isolation, Preparation and Sequencing

RNA was isolated from cells using TriZol (Thermo Fisher) and purified using Zymo Direct-Zol RNA miniprep kit. DNase incubation was carried out for 15 minutes and each sample was quantified using a NanoDrop (Thermo Fisher). cDNA was generated from 1µg RNA using iScript cDNA Synthesis Kit (BioRad). Relative mRNA levels were determined using qPCR reactions performed on the QuantStudio™ 6 Flex Real-Time PCR System with TaqMan™ Fast Advanced Master Mix (Thermo Fisher Scientific) and taqman probes designed to Cdkn1a (Mm04205640_g1, Hs00355782_m1), Mdm2 (Mm01233136_m1, Hs00540450_s1), p53 1-2 (Mm01731287_m1), p53 5-6 (Mm01731290_g1), p53 8-9 (Mm00441964_g1), Bbc3 (Mm00519268_m1), Gadd45a (Hs00169255_m1), 14-3-3σ (Hs00968567_s1). Levels of mRNAs were normalized to the level of Gapdh mRNA. Relative mRNA levels were determined from experimentally determined standard curve (Applied Biosystems). Experiments were carried out in technical triplicates and analyzed using ANOVA with Tukey Post-Hoc Analysis.

Preparation of RNA library and 150bp paired end transcriptome sequencing was conducted by Novogene Co., LTD (Beijing, China). STAR and Tophat alignments were performed with default parameters to generate RNA-seq BAM files with the Genome Reference Consortium Mouse Build 39 genome (GRCm39)(46,47). Indexing with samtools and sashimi visualization was done in Integrative Genomics Viewer. Stringtie novel transcript discovery was carried out using default parameters and plotted using ballgown suite in R.(48)

### Incucyte Growth Tracking

100 cells were plated in a 96 well plate and placed in an incubator outfitted with the Sartorius Incucyte Live cell analyzer. Cells were plated in 2µg/mL doxycycline containing media or control media for six days and analyzed using companion Incucyte software. Experiment completed in ten technical replicates and analyzed by ANOVA.

## Acknowledgments

The authors acknowledge the Duke Cancer Institute Transgenic Mouse Core for their assistance in designing the p53^E221D^ targeting construct and for generating the p53^E221D^ mice. The authors acknowledge the Functional Genomics Shared Resource for their assistance in design and generation of the doxycycline inducible H1299 p53 mutant cell lines. This work was supported by the National Cancer Institute for Dr. David G Kirsch, NIH (R35-CA197616) and National Cancer Institute R03CA249562 and Whitehead Scholar Award from Duke University School of Medicine awarded to Dr. Chang-Lung Lee

## Authors’ Contributions

I.C.L., N.H.L., W.F., and D. G. K. designed experiments. I.C.L., N.H.L., W.F., E.C.M, L. L., and Y. M. performed experiments, I. L., N.H.L. and W. F. performed immunoblots and tissue culture. Y.M. performed histology, and D. M. C. analyzed H&E and IHC slides. I.C.L., E. S. X., L. L. bred and maintained animals. C.L.L and D. G. K. interpreted results and provided critical advice for the manuscript. I.C.L., C.L.L, and D. G. K. drafted the manuscript. All authors edited the manuscript.

Co-Corresponding authorship is requested given the mentorship that Dr. Chung-Lung Lee has provided as Ian seeks to support, revise, and submit this manuscript Dr. David Kirsch provided.

## Data Availability

RNA-Seq data were deposited into the Gene Expression Omnibus database under accession number: GSE239745.

---

**Supplementary Figure 1:**

A. Targeting strategy for CRISPR mediated, homology directed, repair of endogenous p53 gene for generation of p53 P53^E221D/E221D^ embryos

B. In-vitro guide screening of targeting construct

C. Sanger analysis of three founder mice harboring the P53^E221D/E221D^ mutation

**Supplementary Figure 2:**

Note: Due to the constraints of COVID19 procedures, there was a reduced capacity for consistent mouse observation. This resulted in a reduced capacity for collection of recently deceased and a likely bias towards gross morphological tumors reaching humane endpoints and subsequent collection. Therefore, we provide these data as a qualitative indication of a conserved tumor spectrum rather than a quantitative assessment of subtype prevalence.

A. Pie chart of tumor spectrum observed in p53^E221D/E221D^ littermate controls.

B. Pie chart of tumor spectrum observed in p53^E221D/WT^ littermate controls. Mouse 640847 presented multiple primary tumors, one lymphoma and one sarcoma and thus is counted in both categories.

**Supplementary figure 3:**

A. Bar chart comparison of known transcripts and novel splice events in Stringtie results of bam files aligned with Tophat and Star from p53^E221D/E221D^ MEF samples. (Blue = Tophat aligned, Green = Star aligned)

B. Bar chart comparison of known transcripts in Stringtie results from Tophat and Star aligned reads from p53^WT^ MEF samples. (Blue = Tophat aligned, Green = Star aligned)

C. Bar chart comparison of known transcripts and novel splice events in Stringtie results of bam files aligned with Tophat and Star from p53^E224D/E224D^ NCI-H716 samples. Novel transcripts aggregated by the variation at the intron six 5’ splice donor site. (Blue = Tophat aligned, Green = Star aligned)

D. Bar chart comparison of known transcripts in Stringtie results from Tophat and Star aligned reads from p53^WT^ HCT116 samples. (Blue = Tophat aligned, Green = Star aligned)

E. Cryptic splice site in novel transcript, labeled 5bp intron inclusion in Supplementary Figure 2C, called in Stringtie results of Star alignments includes five base pairs of the intron in human cancer harboring the p53^E224D/E224D^

