## Supplementary figures and images for "Mis-splicing Drives Loss of Function of p53^E224D^ Point Mutation"

### Supplemental Figures

#### Supplementary Figure 3.

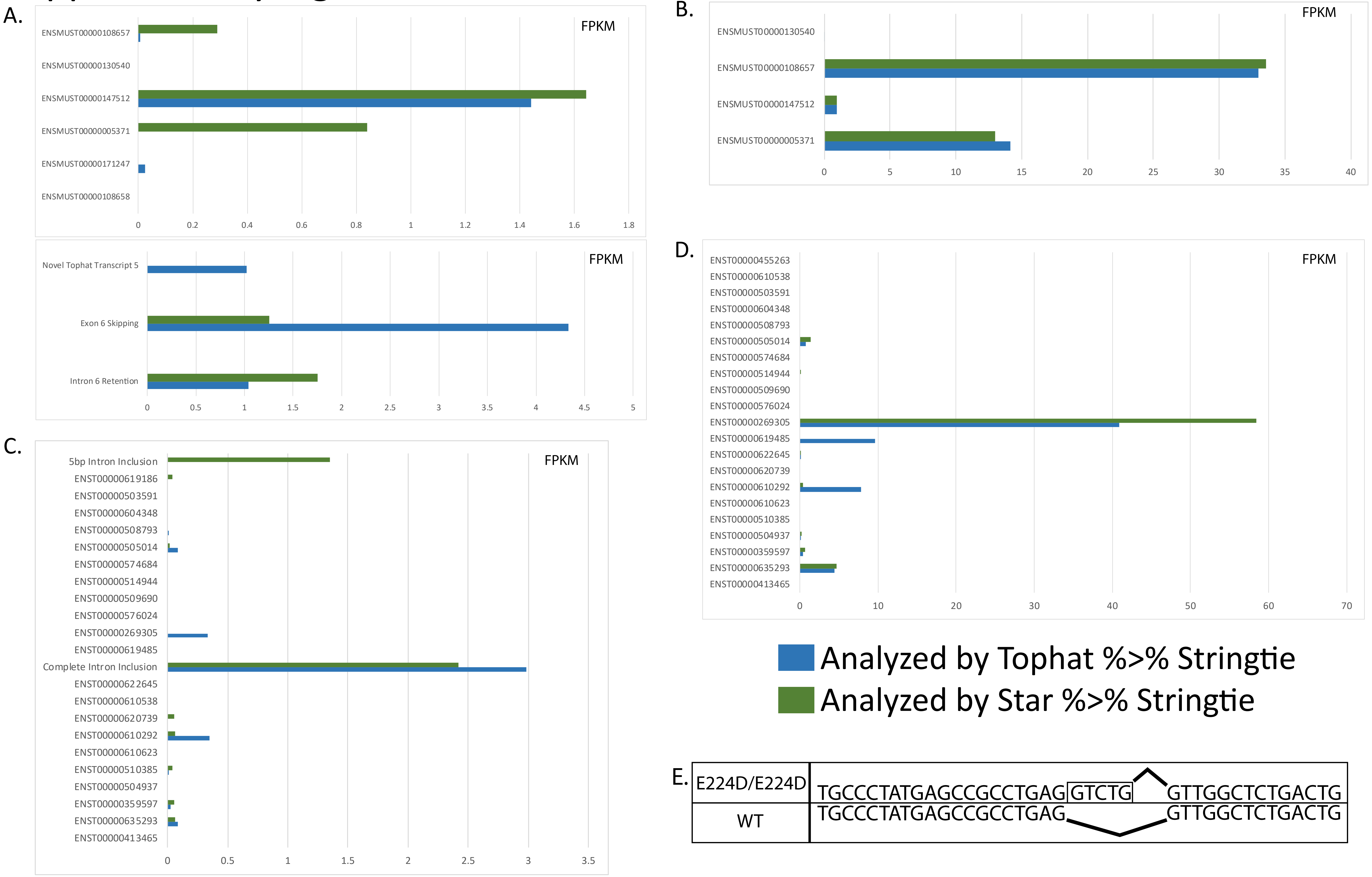
